# What did the dove sing to Pope Gregory? Ancestral melody reconstruction in Gregorian chant using Bayesian phylogenetics

**DOI:** 10.1101/2024.08.19.608653

**Authors:** Gustavo A. Ballen, Klára Hedvika Mühlová, Jan Hajič

## Abstract

An attractive goal in the study of Gregorian chant melodies is reconstructing unobserved melodies as they may have been transmitted along the history of chant, especially as early chant notation does not capture pitch exactly. We propose doing this computationally using Ancestral State Reconstruction (ASR) over phylogenetic trees. Bayesian phylogenetic trees have shown promise as a tool to study the evolution of chant melodies, by inferring a plausible topology of chant transmission. However, the inferred trees cannot be used as ASR inputs directly, because they are undirected, and their branch lengths conflate time and evolutionary rate. We therefore first apply Divergence Time Estimation (DTE) to separate them and represent the tree in a directed form on the time dimension. Using ASR, we then obtain reconstructions of melodies for each of the ancestral nodes, in addition to their distribution in time obtained from DTE, and thus we obtain a phylogeny of chant melody with a music-historical interpretation. We applied this method to the Christmas Vespers dataset, and compare the results against musicological knowledge and melodies reconstructed at Solesmes using methods of contemporary philology, which shows potential for reconstructing cultural transmission through time.

## 1 Introduction

Gregorian chant is the universal liturgical monody of the Roman Catholic church, a major part of Western music history, and a dominant musical tradition up until at least the early 17th century. Its moniker comes from the legend of pope Gregory I., who supposedly dictated Gregorian repertoire as sung to him by a dove of the Holy Spirit. After the 2nd Vatican Council in the 1960s, the role of chant diminished, but it remains highly standardized, with a global authoritative edition: today, apps exist that contain the texts and melodies to be sung at each point of the day’s liturgy (e.g., ChantTools, SquareNote, and Neumz)

However, in the manuscript tradition between approx. 800 and 1600, one rarely finds the melody of a chant written exactly the same in two different sources. Exact pitch notation spread only after 1050 AD, which is also the case for melodies encoded in the adiastematic neume system. The Gregorian melodies spanning all of Latin Europe for the better part of a millenium, were – despite the institutional conservation pressure of their sacred nature – diverse. There are long-open questions in chant scholarship about the nature of this diversity [1, 2, 3, 4], supported by the Cantus database, Cantus Index [5], and its digital ecosystem (i.e., https://dact-chant.ca/, SIMSSA Cantus Ultimus https://cantus.simssa.ca/)

We consider the evolutionary perspective on chant, with a root and a tree of “common ancesotrs”, – despite the role of lateral transmission in cultural evolution [6] – justifiable. The Gregorian tradition does follow from a significant transmission event when a Roman tradition was brought and disseminated across the Frankish empire after 754 [7, p.514–518], and the subsequent Gregorian chant manuscript culture strongly relied on memory [8, 9] and copying [10, 11]. Recently, the approach has shown promise by inferring a plausible Bayesian phylogeny of chant melody [12], but the method stops short of providing a music-historical interpretation of the resulting phylogeny, limiting its value to chant scholarship.

We propose an extended pipeline for inferring from chant melodies a phylogeny with a more defined music-historical interpretation, and thus the value of its predictions can be evaluated more accurately. In contrast to the unrooted topology of [12], our method outputs a rooted tree ordered along the temporal dimension, so that every inferred internal node is associated with a distribution in time, which in turn enables sampling ancestral melodies from the posterior. We build this enriched Bayesian phylogeny of the Christmas Eve vespers dataset. With a lack of known ground truth to compare the resulting phylogeny to, we must instead evaluate the result against musicological knowledge, including the 20th-century Solesmes editions of chant, which (partially) aimed to reconstruct “original” chant melodies that, according to legend, the dove whispered to St. Gregory.

In 2023, Bayesian phylogeny inference using the sequence of pitches in a set of aligned melodies as data has been proposed [12], which sampled the posterior distributions of the unrooted topology as well as branch lengths, which were unfortunately not easy to interpret in a music-historical context. This limitation has directly inspired our present work. There are works from the history of chant scholarship that show evidence for the existence of certain melodic dialects [3, 1], which should in fact be expected due to the oral nature of chant transmission [13, 14, 11], even though the extent of this orality is unclear [15].

The evolutionary perspective on cultural evolution has been applied in music [16] as well as in other areas, for instance in linguistics [17], and human tool development such as knots [18] and palaeolithic artifacts [19]. A phylogeny of Gabon folk music patrimonies has been proposed [20], which highlighted the role of vertical transmission, compared to the focus on lateral transmission in evolutionary linguistics [6]. Bioinformatics tools have been used to characterize diversity in Japanese vs. English and US folk melodies directly [21], or electronic music [22]; most recently, Billboard songs have been studied using methods from evolutionary cancer genomics reformulated as a variational autoencoder [23]. We also must mention the Cantometrics project [24], which proposed a high-level evolutionary tree of folk music worldwide. This ethnomusicological perspective can justifiedly be applied to chant [25].

With the exception of [21], these works extract features from their data that are known from other literature to be salient for the research questions posed. However, for chant melody (as opposed to features related to tonality for later European music), such features are not known yet. Computational work on chant melody has so far tried segmentation [26, 27, 28, 29, 30], but despite indications of the formulaic nature of parts of chant repertoire [31, 32], these experiments have not yet led to a satisfactory theory of chant melody. Thus, we must use the chant melodies directly, with the attendant requirement of directly comparable melodies from different sources.

From the overview of related work, it is clear that computational chant scholarship, is gaining momentum. However, the gap between computational results and their music-historical interpretation remains wide [12, 30]. This work aims to provide a method that significantly reduces this gap with predictions directly testable against concrete musicological knowledge.

## 2 Materials and Methods

### 2.1 Data: The Christmas Eve Vespers and the Solesmes Edition

The dataset of Christmas Eve vespers, used in [12], is so far the best that we have available with a set of fully transcribed directly comparable melodies from a diverse selection of 14 sources: two secular French, three Cistercian, three Benedictine, one Augustinian, a secular source from the Low Countries, and a set of later secular sources from Bohemia. The dataset was originally collected in order to study relationships between late medieval Bohemian sources with the data including transcribed melodies available in the Hymnologica database (http://hymnologica.cz/jistebnice), and we combined this data with all further melodies available for Vig. Nat. Domini vespers from the Cantus Index interface, in order to cover a broader European context. Because the repertoire in office sources is not entirely consistent across sources and our system aligns melodies directly, we had to select a subset of chants contained in as many sources as possible, and then reduce the set of sources to those that contained as many of these chants as possible. The resulting dataset contained 14 sources, with fully transcribed melodies for the following Cantus IDs: 001737 (*Orietur sicut sol*, antiphon), 002000 (*Cum esset desponsata*, antiphon for the Magnificat), 003511 (*Judaea et Jerusalem*, antiphon), 004195 (*Bethleem non es minima*, antiphon), 007040a (*Constantes estote videbitis*, responsory verse), and 007040 (*Judaea et Jerusalem*, responsory). The responsory Judaea et Jerusalem has previously been misreported as Cantus ID 605019, which refers to a shorter version of the reponsory which uses the text “*Cras egrediemini*…” as a verse instead of respond, while 007040 uses “*Constantes estote*…” as the verse (Cantus ID 007040a). A full overview is given in Tab. 1.

**Table 1:** Sources of the Christmas Vespers dataset with their provenance, approximate date, cursus, and presence of the chant in each source (1 or more instances per source). NA represents chants not present in a given source.

| Source | Provenance | Date | Cursus | 605019 | 001737 | 002000 | 003511 | 004195 | 007040a |
| --- | --- | --- | --- | --- | --- | --- | --- | --- | --- |
| A-Wn 1799** | Rein | 1200s | Cistercian | 1 | NA | 1 | 1 | 1 | 1 |
| A-VOR Cod. 259/I | Prague | 1360 | Secular | 1 | 2 | 1 | 1 | 1 | 1 |
| CDN-Hsmu M2149.L4 | Salzinnes | 1554 | Cistercian | 1 | NA | 1 | 1 | 1 | 1 |
| CH-E 611 | Einsiedeln | 1300s | Benedictine | 1 | 3 | 1 | 1 | 1 | 1 |
| CZ-HKm II A 4 | Hr. Král. | 1400s | Secular | 1 | 1 | 1 | 1 | 1 | 1 |
| CZ-PLm 504 C 004 | Pilsen | 1616 | Secular | 1 | 1 | 1 | 1 | 1 | 1 |
| CZ-Pu XVII E 1 | Bohemia | 1516 | Unknown | 1 | NA | 1 | 1 | NA | 1 |
| CZ-Pn XV A 10 | Prague | 1300s | Secular | 1 | 1 | 1 | 1 | 1 | 1 |
| CZ-Pu I D 20 | Passau | 1300s | Augustinian | 1 | 1 | 1 | 1 | 1 | 1 |
| D-KA Aug. LX | Zwiefalten | 1100s | Benedictine | 1 | 1 | 1 | 1 | 1 | 1 |
| D-KNd 1161 | Köln | 1200s | Cistercian | 1 | NA | 1 | 1 | 1 | 1 |
| F-Pn lat. 12044 | Paris | 1100s | Benedictine | 1 | 1 | 1 | 1 | 2 | 1 |
| F-Pn lat. 15181 | Paris | 1300s | Secular | 1 | NA | 1 | 1 | 2 | 1 |
| NL-Uu 406 | Utrecht | 1150 | Secular | 1 | 2 | 1 | 3 | 2 | 1 |

We originally intended to use the CantusCorpus dataset [26] of Office melodies. However, the authors of the Cantus Database preferred transcribing entire sources, so while there are more than 13000 fully transcribed antiphons in CantusCorpus v0.2 [27], the vast majority comes from less than 20 sources. This is further compounded by the surprising diversity of office repertoire. Thus, in the entirety of CantusCorpus, it is only possible to find 10 different sources that have transcribed melodies for the 5 antiphons. Hence, we decided to use the Christmas dataset, with its advantage of having been collected specifically in order to make the comparison between different sources possible.

For each of the 14 sources, we briefly report its century of origin, its provenance, and which ecclesiastical institution it belonged to.

- **A-VOR Cod. 259/I**. A 14th century antiphoner of the collegiate chapter church of Vyšehrad, Prague. In the early 15th century, it was moved to Vorau because of Hussite wars. In 1490-1500, it was adapted for Salzburg liturgy. Available at https://manuscripta.at/hs_detail.php?ID=6267.
- **A-Wn 1799\*\***. A 13th century Cistercian antiphoner from the Rein monastery in Austria. Available at https://differentiaedatabase.ca/manuscripts/wn-1799.
- **CDN-Hsmu M2149.L4**. Cistercian antiphoner from the Abbey of Salzinnes, Namur, in the Diocese of Liège, central Belgium, completed in 1554-1555. Available at https://cantus.uwaterloo.ca/source/123723.
- **CH-E 611**. A 14th-century antiphoner from the Benedictine monastery of Einsiedeln, central Switzerland. Available at https://cantus.simssa.ca/manuscript/123606/.
- **CZ-HKm II A 4**. An antiphoner from the 1470s, belonging to the municipal Church of the Holy Spirit in Hradec Králové, eastern Czechia. Available at http://hun-chant.eu/source/1481?page=1.
- **CZ-PLm 504 C 004**. A late antiphonary from the St. Bartholomew municipal church in Pilsen, western Czechia, from 1616. Available at https://rukopisy.zcm.cz/view.php?ID=504C004.
- **CZ-Pn XV A 10**. Late 15th century notated breviary from the cathedral cursus in Prague, Czechia. Available at http://hymnologica.cz/source/47.
- **CZ-Pu I D 20**. An antiphonary from the Augustinian monastery in Třeboň, southern Czechia, created in the 2nd half of the 14th century. Available at http://hymnologica.cz/source/10721.
- **CZ-Pu XVII E 1**. A mixed Latin and Czech antiphonary from the early 16th century, of Czech (but further unspecified) provenance. Available at http://hymnologica.cz/source/10664.
- **D-KA Aug. LX**. A complex 12th-century antiphoner, of which the musical notation was almost completely rewritten in the 13th or 14th centuries. From the Zwiefalten Benedictine monastery in southwestern Germany, moved to the abbey of Reichenau in the 15th century. Available at https://cantus.uwaterloo.ca/source/123612.
- **D-KNd 1161**. A late 12th- and early 13th-century Cistercian antiphoner, possibly written for use by the female abbey of Saint Mechtern in Cologne, western Germany, renamed Saint Apern in 1477. Available at https://cantus.uwaterloo.ca/source/601861.
- **F-Pn lat. 12044**. An early 12th-century antiphoner from the Benedictine abbey of St.-Maur-de-Fossés, close to Paris, France. Available at https://cantus.uwaterloo.ca/source/123628.
- **F-Pn lat. 15181**. An early 14th-century notated breviary belonging to the Notre Dame cathedral in Paris, France. Available at https://cantus.uwaterloo.ca/source/123631.
- **NL-Uu 406**. A 12th-century antiphonary from St. Mary’s church in Utrecht, Netherlands. Later 13th-15th-century changes. Complex source that has multiple versions of some melodies. Available at https://cantus.uwaterloo.ca/source/123641.

The three major dimensions of external similarity between chant sources, in terms of how similar the segments of culture represented in these sources are expected to be, are geography, chronology, and *cursus*, i.e., space, time, and the liturgical context within which the books were used. It is not entirely clear in chant scholarship how strongly each of these factors should influence chant melodies (the exception where *cursus* is clearly expected to dominate other factors is that of the Cistercian order, which mandated that all monasteries must have identical liturgical books [33, p. 99]), but these organizing principles should be observed in the resulting tree.

Solesmes editions are found in the Gregobase corpus (https://gregobase.selapa.net), from where we collected the melodies corresponding to our inference dataset. However, only the responsory *Judaea et Jerusalem*, its verse *Constantes estote videbitis*, and the antiphons *Judaea et Jerusalem, Orietur sicut sol*, and *Cum esset desponsata* are available.

#### 2.1.1 Preprocessing

Some sources contained multiple instances of chants of one Cantus ID: In that case, we retained the version with the most complete version of the melody (as repeated instances of the same chant are sometimes only written as incipits in the sources), and if multiple full melodies were available, we selected the melody that was directly in the Vig. Nat. Dom. section (Table 1).

Furthermore, as noted by [30], differentiae (see https://differentiaedatabase.ca/about for details) tend to be encoded together with antiphon melodies and must be removed. Furthermore, in the Cistercian sources, some melodies had been recorded transposed a fifth up. This was a scribal practice based in the Cistercian reform of chant theory which has no implication on performance [34], so we transpose these melodies back by a fifth down, in order to not introduce spurious differences.

### 2.2 Methods

We extended an earlier pipeline [12] that aimed to reconstruct the unrooted phylogenetic tree of sources by adding three additional steps: Model selection for the root using Bayes factors, divergence time estimation on the fixed rooted tree selected in the previous step, ancestral melody reconstruction using the resulting timetree, and finally comparison with the melodies reconstructed in the Solesmes edition of chant (Figure 1).

**Figure 1:**
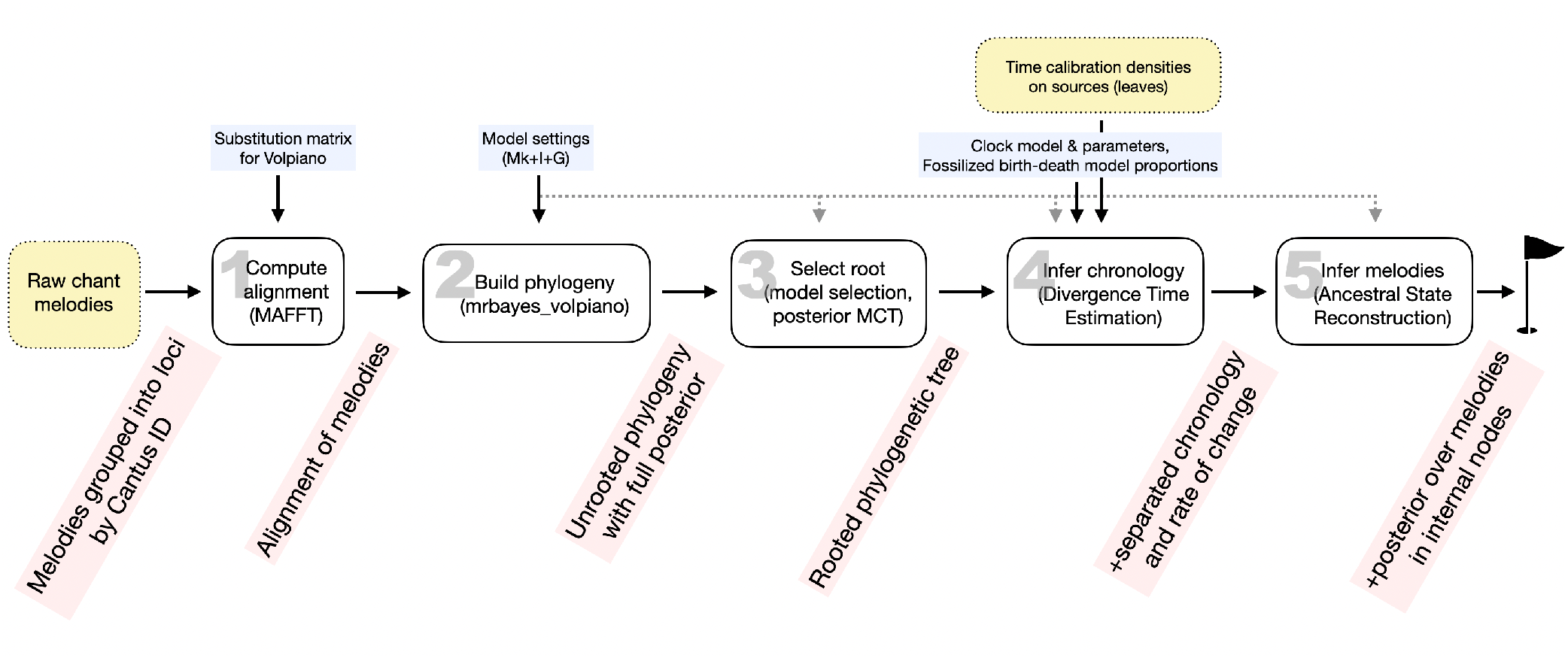
Overview of the whole pipeline, from Multiple Sequence Alignment to melody reconstructon. Externally provided data is highlighted in yellow, parameters in blue, artefacts at intermediate steps in red. Steps 1 and 2 have been described in [12], in this paper we focus on steps 3-5. These steps start from an unrooted phylogeny, which – while its topology already provides some insight into the development of chant melody – has no chronology and therefore no music-historical interpretation; and they lead to a phylogeny that assigns to all its inferred internal nodes a distribution in time and a posterior distribution over the melodies corresponding to that node, therefore making predictions that can then be verified against music-historical knowledge, and provide new insights as well.

A Bayesian phylogeny [35] of chant melodies was carried out following the approach by [12] using `mrbayes_volpiano` [36, 12] (available at https://github.com/gaballench/mrbayes_volpiano). The resulting tree is unrooted and has limited interpretation in terms of cultural transmission, because it is not clear how time flows along the tree (as opposed to rooted phylogenetic trees). Also, branch lengths in unrooted trees are measured in units that conflate evolutionary rate and time [37]. We identify rate and time using more sophisticated methods [38] to measure evolutionary rate and its heterogeneity along the tree, and also to use time as the unit of branch lengths [39]. At that point we chose a single tree on which to carry out further analysis, and then calculated a maximum clade credibility tree as a summary tree from the posterior distribution of topologies, for which we used `treeannotator` [40]. Tree formatting was carried out with `phyx` [41].

#### Model selection for the position of the root

There is no obvious way to root a Gregorian chant phylogeny as it appeared as a unique event in time with no sisters that we can explicitly use in phylogenetic analysis, at least not when one considers notated sources with comparable repertoire. A good choice would be some peripheral liturgy, but unfortunately the closest such candidate – Ambrosian/Old Roman chant – does not have comparable repertoire for the liturgical positions available in our dataset (and in any case its evolutionary relationship to Gregorian repertoire is unclear) [7, pp. 530-539].

Because we lack any indication of an outgroup with sequence data for analysis we can consider the root position as a fixed model and then calculate the marginal loglikelihood (*lnL*) using stepping stones under a Bayesian framework [42]. This approach allows model selection using Bayes factors 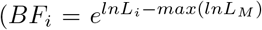 where *i* is a given model out of *M* alternatives) while keeping the age information, the data, and other model parameters identical across analyses. An unrooted binary tree with *N* terminals has 2*N* − 3 branches, and therefore we potentially can root on each of them as the varying part of the model on these *M* = 2*N* − 3 model settings. Once picking the best root position via BF and model posterior probability 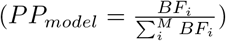, we can use the best rooted topology to carry out DTE.

Generation of all possible rooted trees on the MCC tree was carried out using the packages `ape` [43] and `phytools` [44] in `R` v.4.3 [45]. We applied stepping stones in `mrbayes_volpiano` with 50 stones and 1.000.000 generations sampling every 100th generation. The marginal likelihood was then used for calculating BF and model PP in `R`.

#### Divergence Time Estimation (DTE)

Once we have selected a rooted tree Ψ we can use Bayesian DTE on the fixed topology in order to separate evolutionary rate *r* from time *τ* in branch lengths [46]. The Bayesian model in general form (Equation 1) uses time calibration densities *f* (*τ*) on either the nodes or the tips in order to generate the joint time prior *f* (*τ*, Ψ) using the calibration densities 2 and the tree process. Separation of rate and time requires also a clock model *f* (*r*|*τ, θ*) which describes the branch-specific evolutionary rates. The model parameters *θ* include parameters of the tree process, the clock model, and the evolutionary model. Then, the model estimates the PP densities for all of the model parameters conditional to the melody alignment data *D*.

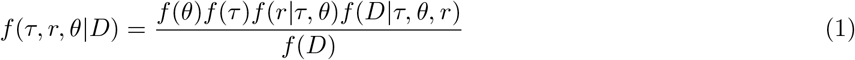

It is possible to extend the Bayesian model by specifying the tree prior using a fossilised birth-death model parametrised using speciation, extinction, and sampling proportion (equation 1 in [47]). These parameters are often difficult to model from previous knowledge, but it is possible to use “uninformative” or flat priors for them [48].

DTE was carried out in `mrbayes_volpiano` using the Mk model with inverse-gamma rates [49]. The rooted tree topology was fixed. The clock model was set to independent gamma rates [50] and an Exponential(2) prior used for the variance whereas an Exponential(5) prior was used for the clock rate. The FBD parameter priors were set to uniform(0,10) for the speciation rate, and Beta(1,1) for both the extinction and fossilisation rates. Calibration densities were specified as in Table S1. Parallel-tempering MCMC [51] was used with eight chains, one cold and seven hot, with a temperature of 0.001. MCMC sampling was carried out with 10.000.000 generations, sampling every 1.000. Posterior summarisation used a burn-in of 10%. All parameters attained effective sample size (ESS) *>* 500, indicating an appropriate sampling of the parameter posterior distributions. Time tree plotting (Figure 3) was carried out in `figtree` [52].

#### Ancestral State Reconstruction (ASR)

These methods infer most likely state of the sequence (position-wise) on the resulting posterior tree. This needs a fully resolved tree, so instead of the summary tree we use the maximum credibility tree. It is possible to generalise the independent ASR for each character in the alignment and then to collect the reconstructed states to represent a whole sequence, or in our case study, a melody, therefore producing an Ancestral Melody Reconstruction (AMR). ASR requires an evolutionary model for discrete states which in this case is the same the one used for phylogenetic inference (i.e., the Mk model).

Now we can use the Mk model for calculating the probability of transition as a function of time [46, 53]. Here time are the branches, and we start from the observed states at the tips and traverse the tree towards the root estimating the ancestral states. Simulation is used to sample form the PP density for each of the states using stochastic mapping [54]. Because we have PP densities for each melody position at each internal node, we can calculate posterior probabilities for arbitrary melodies. For instance, we can take an aligned melody generated by another reconstruction method and calculate the probability of observing it at any arbitrary node of the tree under the PP density of states.

AMR was carried out using the equal-rates model in `phytools` [44]. Stochastic character mapping was used for constructing the posterior distribution on each node using empirical Bayes using 1000 iterations. This process required parallelising the ASR for each position in the melody using the packages `doParallel` [55] and `foreach` [56] in `R`.

The Solesmes melodies (see 3.3) were aligned to the original alignments used for tree inference using `mafft` [57] with the `--keeplength` argument so that the base alignments remain unchanged. We then collected the maximum *a posteriori* states in order to construct the AMR on each node. Finally used the posterior of states on each internal node in order to calculate the per-node PP for both the AMR and the Solesmes reconstruction. Note that when producing MAP estimates of melodies using ASR, sometimes there is a tie at a certain position. This can mostly be resolved by preferring a note over a gap. In the rare case when there is a tie between two notes, we arbitrarily prefer the lower note.

#### The Solesmes edition

The comparative approach of Solesmes means they should not be included in building the phylogeny as the assumption about an evolutionary process does not hold, because such melodies did not evolve naturally through history but represent musicological reconstructions instead. However, because we have for each internal node a posterior distribution over melodies, we can compute the probability of observing the given Solesmes melody on each inferred node. Finally we calculated the joint probability of observing all the Solesmes melodies at a given node.

#### Code and data availability

All the code and data necessary for reproducing the results herein presented are available on GitHub (https://github.com/Genome-of-Melody/divtime_christmas) as well as on Zenodo (DOI https://doi.org/10.5281/zenodo.13344525).

## 3 Results

The topology of the phylogeny on the Christmas dataset follows the same contours of geography and c*ursus* as the phylogeny of [12]. This is expected, as the alignment is similar and the tree inference methods are same. However, as the dataset has been cleaned, there are a few changes in the topology. There is a distinct French and Cistercian branch (given that the Cistercians originated in France, this is not surprising), then a geographical gradient towards the group of secular Bohemain sources (CZ and A-VOR), with a monastic group of Benedictine and Augustinian sources (CZ-Pu I D 20, D-KA Aug. LX, and CH-E 611). The historical Augustinian link between CZ-Pu I D 20 and A-VOR 259 I is, however, not replicated.

**Table 2:** Calibration densities (CD) used in DTE. Time scale is in both years before the present (YBP, as used by mrbayes_volpiano) as well as in anno Domini (AD). Single time values represent fixed values whereas intervals represent Uniform(min,max) calibration densities.

| Node | CD (YBP) | CD (AD) | Ref |
| --- | --- | --- | --- |
| A VOR Cod 259 I | 654 | 1370 | https://manuscripta.at/hs_detail.php?ID=6267 |
| A Wn 1799 | 724–824 | 1200–1300 | https://differentiaedatabase.ca/manuscripts/wn-1799 |
| CDN Hsmu M2149 L4 | 474 | 1550 | https://cantus.uwaterloo.ca/source/123723 |
| CH E 611 | 624–724 | 1300–1400 | https://cantus.simssa.ca/manuscript/123606/ |
| CZ HKm II A 4 | 554 | 1470 | http://hun-chant.eu/source/1481?page=1 |
| CZ PLm 504 C 004 | 408 | 1616 | https://rukopisy.zcm.cz/view.php?ID=504C004 |
| CZ Pn XV A 10 | 624–674 | 1350–1400 | http://hymnologica.cz/source/47 |
| CZ Pu I D 20 | 624–674 | 1350–1400 | http://hymnologica.cz/source/10721 |
| CZ Pu XVII E 1 | 474–524 | 1500–1550 | http://hymnologica.cz/source/10664 |
| D KA Aug LX | 624–924 | 1100–1400 | https://cantus.uwaterloo.ca/source/123612 |
| D KNd 1161 | 799–849 | 1175–1225 | https://cantus.uwaterloo.ca/source/601861 |
| F Pn lat 12044 | 874–924 | 1100–1150 | https://cantus.uwaterloo.ca/source/123628 |
| F Pn lat 15181 | 674–724 | 1300–1350 | https://cantus.uwaterloo.ca/source/123631 |
| NL Uu 406 | 624–924 | 1100–1400 | https://cantus.uwaterloo.ca/source/123641 |
| Root | 1124–1324 | 700–900 | [7] |

### 3.1 Evolution of melodies in a temporal scale: DTE

#### Root position

A single rooted tree was found to be the best one (*lnBF* compared to the next best tree 1.8; model PP = 0.75) and used for subsequent analysis. The model selection recovered the root position so that the highest-level clades correspond to French and Cistercian sources and non-Cistercian sources coming from the eastern side of the river Rhine in the other clade (Figure 2). The Cistercian order was founded at Citeaux, an abbey in France, and the order was strict about maintaining this lineage also for chant books. This corresponds to existing hypotheses about the distinct “west Frankish” and “east Frankish” chant melodic dialects [3].

**Figure 2:**
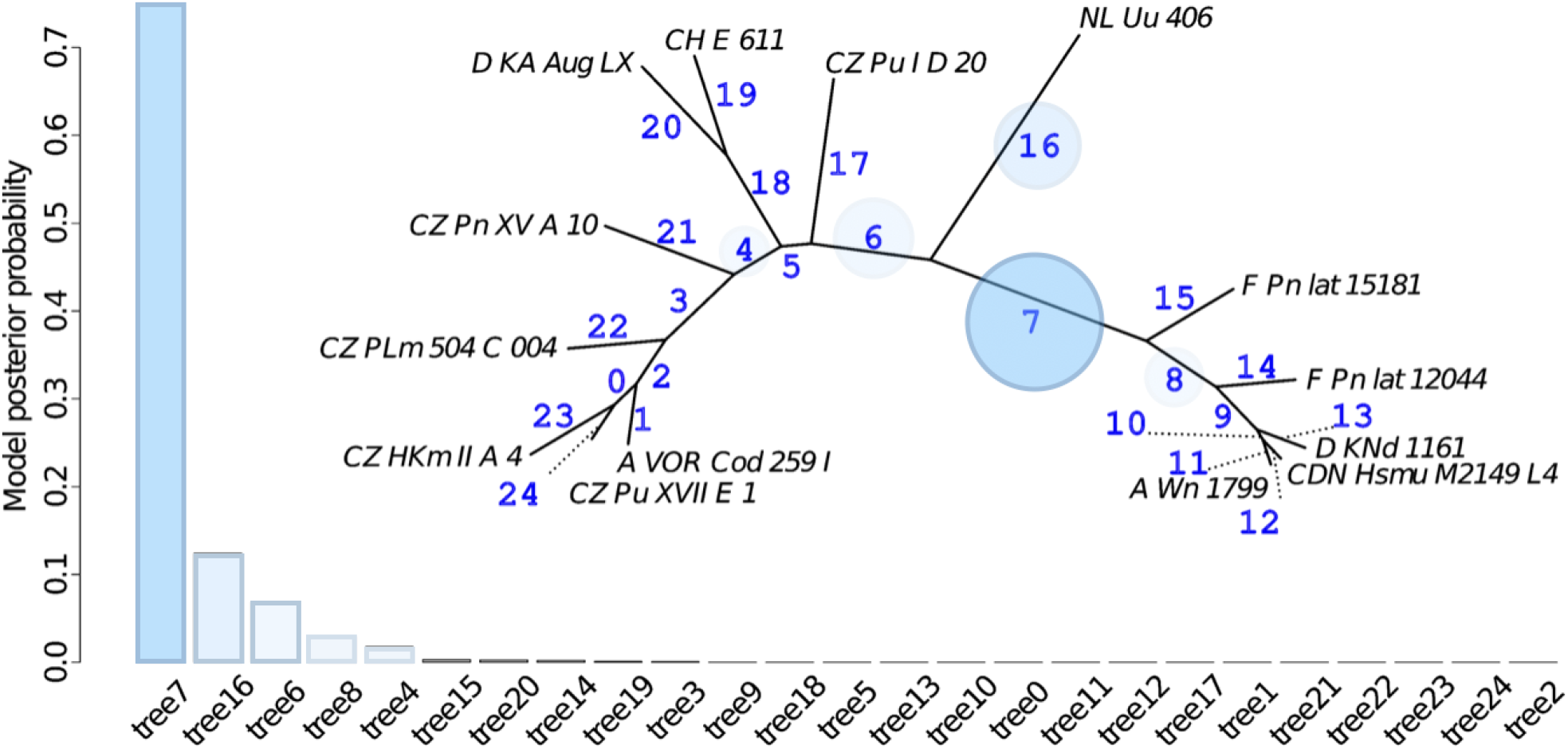
Model selection for the position of the root. Bar height represents posterior model probability, each bar is a topology resulting from rooting at a given branch in the maximum credibility tree. Such trees labeled tree0 through tree24 were the varying element of the Bayesian phylogenetic model for which the marginal likelihood was calculated using the stepping stones method. The best rooting position was found to be the branch labeled as 7 with model posterior probability 0.750, whereas the second best rooting point is the branch labeled as 16 with posterior model probability 0.124.

### 3.2 Estimating ancestral melodies: AMR

Given that the root position found via model selection separates the tree into east Frankish “Germanic” (node 20) vs. west Frankish “Romanic” (node 16) melodies [3, 4], we focus on comparing the melodies inferred for the roots of these two clades. The “Germanic” chant dialect has been characterized by its preference for minor thirds d-f, a-c instead of seconds e-f, a-b (incl. a-b flat) [4], with the insight that it is a general German preference for *re*-*fa* over **re**-*mi* or *mi* -*fa*. A melody in a “Germanic” source exhibiting none this preference merits scrutiny [4, p. 95] (Figure 4).

**Figure 3:**
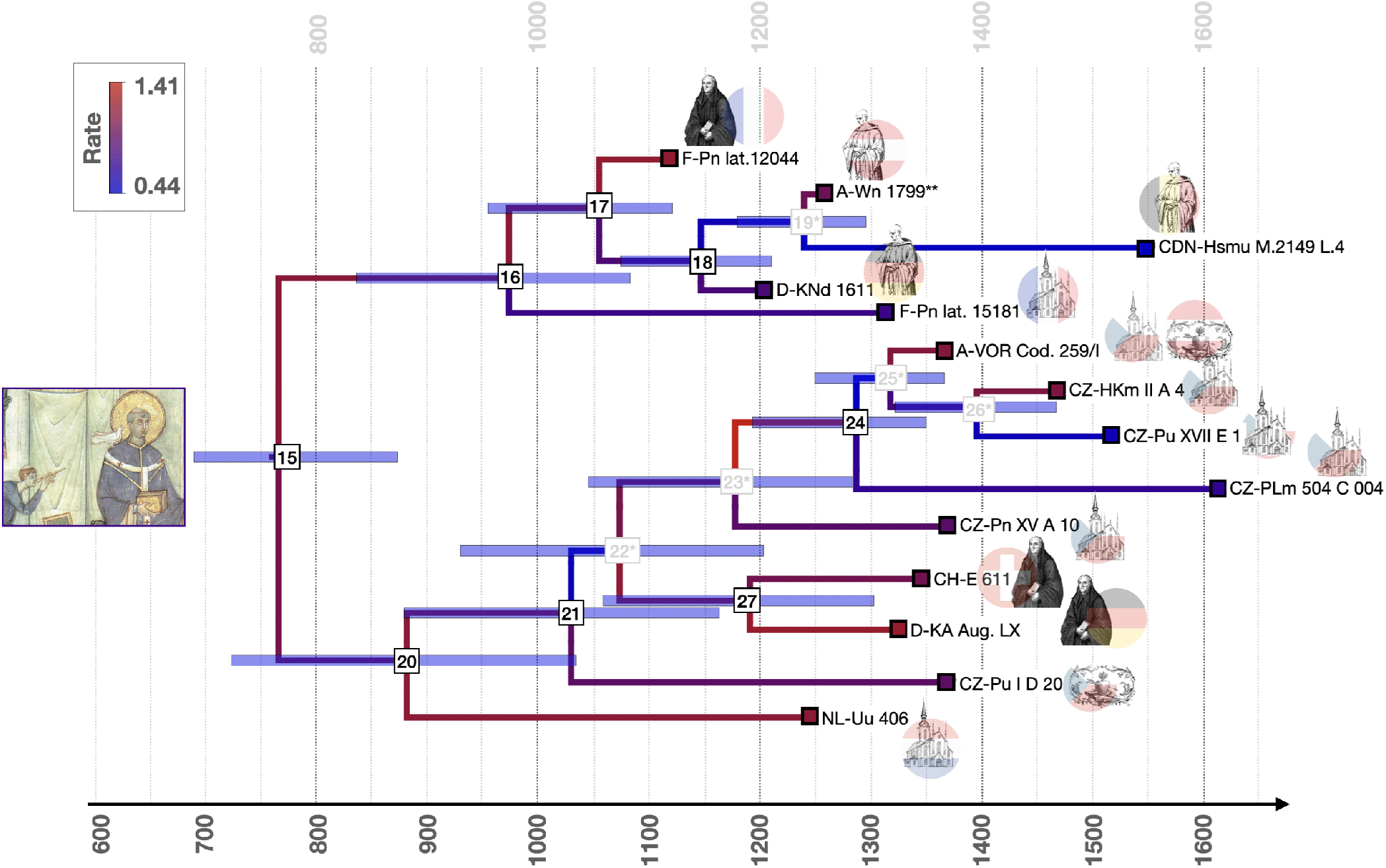
Result of the Divergence Time Estimation algorithm over the Maximum Credibility Tree (MCT) obtained from phylogeny inference and rooted using model selection. The horizontal axis is historical time in years (A.D.), the vertical axis has no meaning. The internal nodes (numbered squares) are positioned at the median estimate of the year they are from. Their numbers are given so that the reported ASR results can be placed on the tree. The blue horizontal bars at internal nodes signify the 95% credibility intervals for the year to which the node corresponds. The color of the branches corresponds to evolutionary rate, with greater expected numbers of changes per year being red. Tips (colored squares) correspond to sources from the Christmas dataset, identified by their sigla. Their region of origin is encoded by a rondel corresponding to the current country governing said location, their cursus by the image (secular – church building, Benedictine – black monk, Cistercian – white monk, Augustinian – heart). The root separates a distinctly French and Cistercian clade (the order was founded in France) from sources east of the Rhine, following the insight of [3]. The grey internal nodes with asterisks are not in the summary tree, signifying that only the topmost common ancestor from the merged set is likely to actually be a common ancestor, and that the down-tree nodes are rather artifacts of requiring the MCT for DTE and ASR computation. Finally, a depiction of the Gregorian legend of the dove dictating melodies to Gregory I. from the Registrum Gregorii (Trier, Stadtbibliothek, Hs. 171/1626) is centered at the time of his papacy (590-604 AD), clearly out of the highest posterior density interval for the root age.

**Figure 4:**
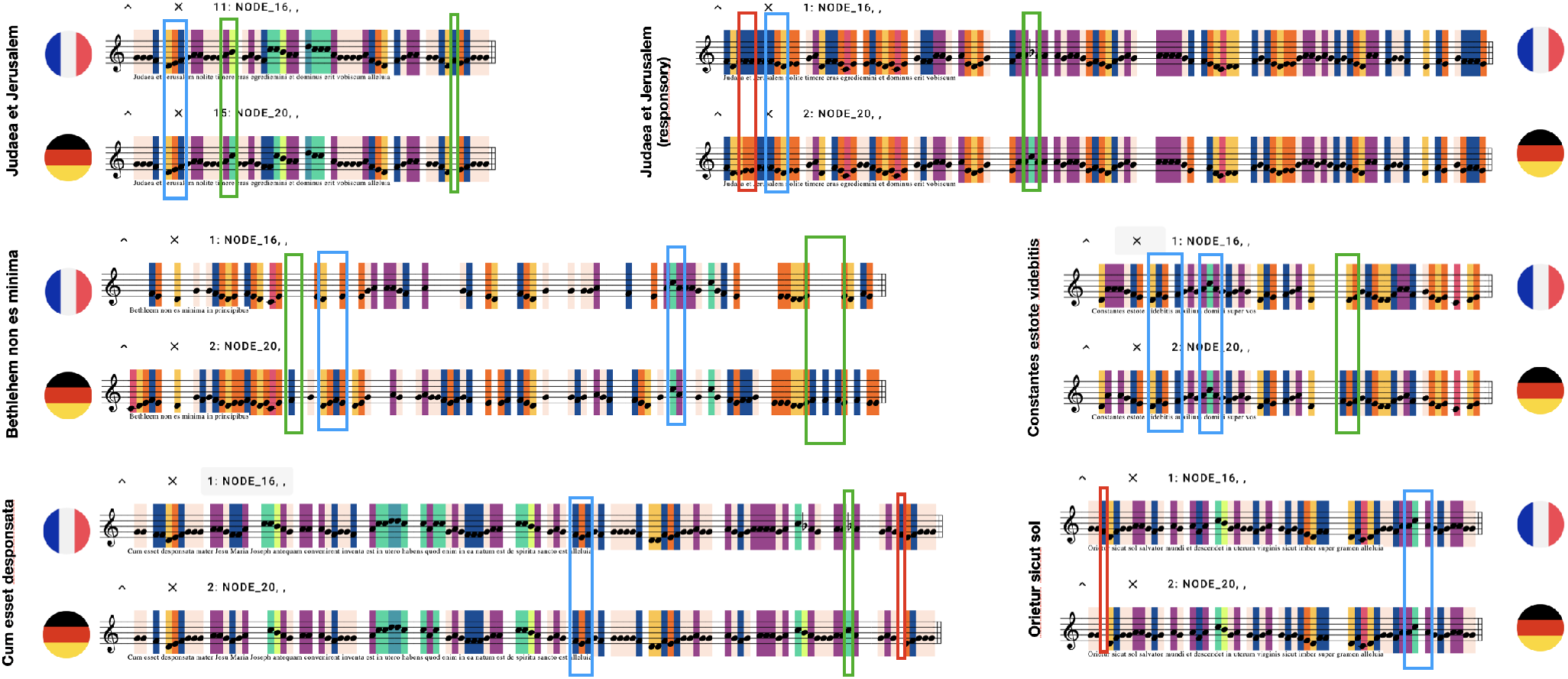
Comparing the results of Ancestral State Reconstruction for the nodes corresponding to the common ancestors of the East Frankish (“Germanic”, node 20) and West Frankish (“French”, node 16) melodic dialects. Each pitch is assigned a background color to make differences in melodies easier to spot. In previous literature on the topic (Peter Wagner’s book “Gregorianische Formenlehre”, 1911), the “Germanic” dialect was supposed to be characterized by its preference of *fa* over the “French” dialect’s preference for *mi*, which corresponds to the “Germanic” dialect preferring the tone *f* over *e* and *c* over *b* (natural, also termed “h” in German nomenclature, as well as flat), which given the prevalence of stepwise motion often translates to preferring thirds *d-f* and *a-c* over seconds *d-e, e-f*, and *a-b(h), b(h)-c*. While the sources weroe “sorted” into these two main melodic dialects by the phylogeny, this preference seems to very weak in the inferred ancestral melodies for these two dialects: while in 7 positions the East Frankish dialect does exhibit this preference (**green** frames), in 3 positions this is the other way around (**red** frames), and most importantly, there are many places where the East vs. West Frankish preferences could have been expressed but is not (some examples in **blue**). More work therefore must be done to uncover what actually differentiates melodies of these two major dialects — whether these are really general preferences, or whether salient preferences are only exhibited in certain parts of melodies. However, in keeping with the observations of Wagner, in each melody there is at least one position where the “Germanic” preference is present.

This is observed in the inferred melodies to a limited extent. For five out of six Cantus IDs in the dataset, at least one such position where node 20 preferred what is in [3, 4] considered Germanic. At the same time, four melodies also have positions where the “Romanic” clade root MAP estimate has “Germanic” features. Out of a total of 77 manually identified “opportunities” for the melodies to differ in their treatment of *re*-*fa* thirds, the “Germanic” option is only chosen by node 20 melody 5 more times than by the node 16 melody. This may partly be an artifact of mode. Most of the Christmas melodies are in mode 8, while the most characteristic examples in [3] and [4] are in mode 1. However, the literature does also indicate that the melodic dialect doesn’t at all manifest at *every* opportunity, and that one might easily find the “Germanic” preference in sources west of the Rhine and vice versa [4, pp.95-98]. Therefore, while ASR results are not as clear a success as accurately predicting the timing of the Cistercian reformation, it does again conform to existing expectation.

### 3.3 Comparing AMR to the Solesmes Editions

The posterior log-probabilities of observing the Solesmes melodies at individual nodes are shown in Fig. 5. At least for the Christmas Eve vespers, the model indicates the editors of Solesmes have preferred melodies of the west Frankish style (nodes 16-19), most prominently those of the Cistercian order (nodes 18 and 19). The melodic features of later secular Bohemian sources (nodes 24-26) are the least preferred models overall, which is mostly influenced by the longer responsory *Judaea et Jerusalem* (judjer2) and the Magnificat antiphon *Cum esset desponsata* (cumesset), not so much by the psalm antiphon *Orietur sicut sol* (orisic) or the responsory verse *Constantes estote* (consest).

**Figure 5:**
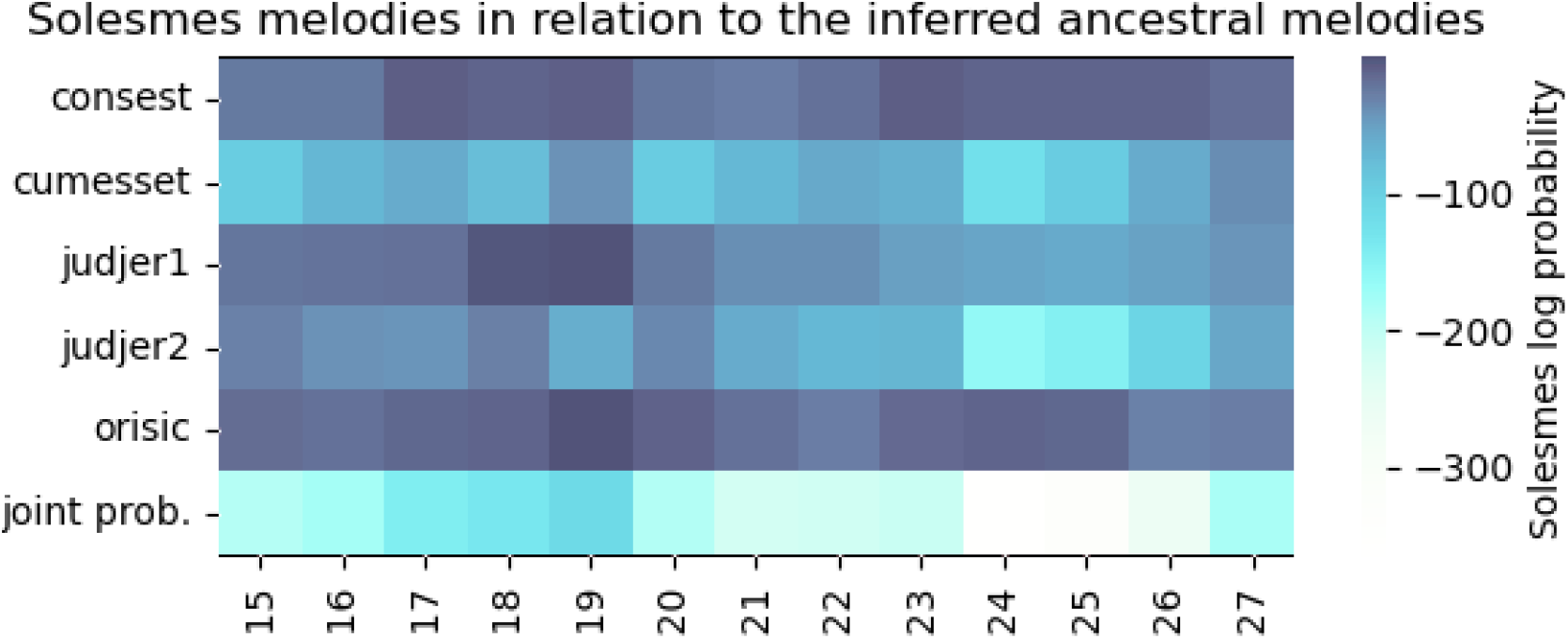
The log-probability of observing individual Solesmes melodies according to the posteriors inferred for internal nodes of the inferred phylogeny, and their joint probability. The node labels are taken from Fig. 3: nodes 17, 18 and 19 correspond to the French and Cistercian monastic melodies, and the nodes with lowest log-probabilities are the Bohemian nodes. Melody names are *Constantes estote videbitis* (consest), *Cum esset desponsata* (cumesset), the antiphon *Judaea et Jerusalem* (judjer1), the responsory *Judaea et Jerusalem* (judjer2), and *Orietur sicut sol* (orisic).

This complements what we know about the process, as well as expectation of the time [1, pp.5-6]. The Solesmes editorial process did not, for unexplained reasons, consider German sources [4, p.113]. Furthermore, the second and third criteria – legitimate tradition and needs of contemporary liturgy, which included the skill sets and habits of early 20th century singers [58, 59] – would gravitate towards a later aesthetic of chant that is closer to the world of tonality, and the Cistercian reform is a significant stepping stone in this respect [34].

## 4 Discussion

We introduce a method for building interpretable phylogenies of chant melody. Because there is no known “true” phylogeny of chant, we rely on comparing the resulting phylogeny to music-historical knowledge. The experiments on the cleaned Christmas dataset and with the Solesmes melodies have provided a number of predictions that could be directly verified against existing knowledge (musicological, codicological, paleographical, historical) and, in this confrontation, have held. The accuracy of the estimated median age of the common ancestor of Cistercian sources is notable. We believe that we have significantly narrowed the gap between computational methods and musicological insight, and that this method – especially when applied to larger datasets, or datasets designed explicitly around open problems – can meaningfully complement philological methods of chant research.

### Cistercians sources

There are three sources in the Christmas dataset of Cistercian provenance: A-Wn 1799**, D-KNd 1611, and CDN-Hsmu M2149.L4. The Cistercian order is known to have maintained very strict discipline in copying liturgical books. This is already seen in the topology inferred in [12], where these sources are grouped closely together, and this is replicated in our topology as well. However, additionally, the DTE step of our pipeline provides an explicit time for the common ancestor of Cistercian sources: the median falls on the year 1145, which coincides with the reform of the antiphonary initiated by St. Bernard of Clairvaux in 1147 or prior [60]. Drafts of the Bernardine liturgy are found in the 12A-B Westmalle Antiphonary, dating from 1140-1143 [34]. The 95 % credibility interval covers 1075 to 1210.

### The position of A-VOR Cod. 259 I

The source A-VOR Cod. 259 I merits attention. It is a source from the late 14th century that was brought to Vienna and later the Augustinian monastery in Vorau when the Chapter house at Vyšehrad (Prague) was fleeing the Hussite wars of 1420-1434, as evidenced by the inscription on its f.1r. This inscription, which describes the history of the manuscript, is written over a rubbed-out earlier melody (Gaude et laetare, Cantus ID 002922, which is often the first antiphon in numerous sources of Bohemian provenance), which is how the new owners found space for the inscription on the first page. Why is this source not grouped closer with either the Augustinian CZ-Pn I D 20, or the similarly late 14th century CZ-Pu XV A 10? In 1496, the manuscript underwent significant revisions according to the Salzburg diocese rite [61]. A look at its digitized folios 69r-71v reveals the presence of possible palimpsests (instances of an older layer scratched out: see https://www.cantusplanus.at/common/rism.php?rism=A-VOR259_1). Some are clearly visible (the doxology at the top of f.71r), and some are just suspicion (traces of rhombic shapes that resemble bleedthrough in the last two staffs on f.69r but do not have corresponding rhombes on f.69v or the neighboring f.68v, which incidentally also has modifications in its bottom half). Furthermore, the antiphon “Gaude et laetare”, overwritten on f.1r by the inscription with the source’s history, is used again on f.70r for Christmas Eve vespers, but with a different melody than what was on f.1r – another reason to suspect that the melodies for Christmas Eve vespers may have been changed later in the 15th century. Of course, codicological expertise is needed to resolve the extent to which this was done; however, this shows how the inferred phylogeny can, aside from providing verifiable predictions such as the Cistercian common ancestor date, also point towards specific problems requiring further musicological study.

If we do accept the later origin of A-VOR Cod. 259 melodies, the branch leading down to node 24 that exhibits the highest evolutionary rate can be symptomatic of two phenomena: one geographic, the Bohemian Reformation, and one chronological – the transition into the late middle ages. The fact that the A-VOR melodies were modified outside of the lands controlled by the Hussites (or, later, Utraquists) lends, in our view, more credence to the chronological explanation (which then leads to unexplored questions about chant melody between the late middle ages and the Editio Medicaea of 1604), but weighing these two interpretations is not possible with the Christmas dataset.

The case study of A-VOR Cod. 259/I shows that this method also brings value in reexamining assumptions about the input data, and thus can also serve as a “quality control” tool. Its improved interpretability further makes it more accessible for musicologists without a significant technical background.

### The Solesmes edition and AMR

We now use our estimated model of chant evolution to study the modern chant restoration movement through the properties of the melodies from the group of 20th- and 21st-century editions of chant produced primarily by the Benedictine monks of the Solesmes Abbey. The Solesmes editions are the central output of the chant restoration movement, led by the Benedictine monks of Solesmes since the latter half of the 19th century [62]. Their core output is the official new edition of Gregorian chant – originally under the auspices of the Editio Vaticana, led by Dom. Pothier and an editorial council, based on the document *Motu Proprio* of Nov 22nd, 1903, of pope Pius X; later editorial activity has been carried out by the monks of Solesmes themselves.

The main contribution of this edition was, in line with the philological thinking of the time, to recover as much as possible the “original” melodies of Gregorian chant, trying to recover – in principle – what the dove had whispered into Gregory I.’s ear. Thus, in the context of ASR, it is enticing to use the Solesmes editions as the “best effort” reconstruction of the melodies that should correspond to the root of our tree, and thus different ways of building the pipeline could theoretically be evaluated by how similar their predicted root melodies are to the Solesmes edition – thus matching this ongoing “best effort” reconstruction using philological methods. However, reconstructing the earliest melodies of chant was not the only priority of Editio Vaticana: another was to do this while retaining “legitimate tradition” from later times, and third, to serve the needs of the day. From the *Motu Proprio* of Pius X.: “The Gregorian melodies are to be restored in their completeness and true nature, according to the testimony of the more ancient manuscripts, taking into consideration not only the legitimate tradition of intervening centuries, but also the common practices of present-day liturgy.” Transl. from [62, p.xxi]. The clashes between these priorities, as manifested in the resultant melodies, were subject to heated exchanges at the time [63, 58, 59]. and the process involved diverse influences such as anticlericalism or business interests of print unions [64]. This can be observed also in the melodies directly when we compare them with the inferred medieval melodies in Fig. 6.

**Figure 6:**
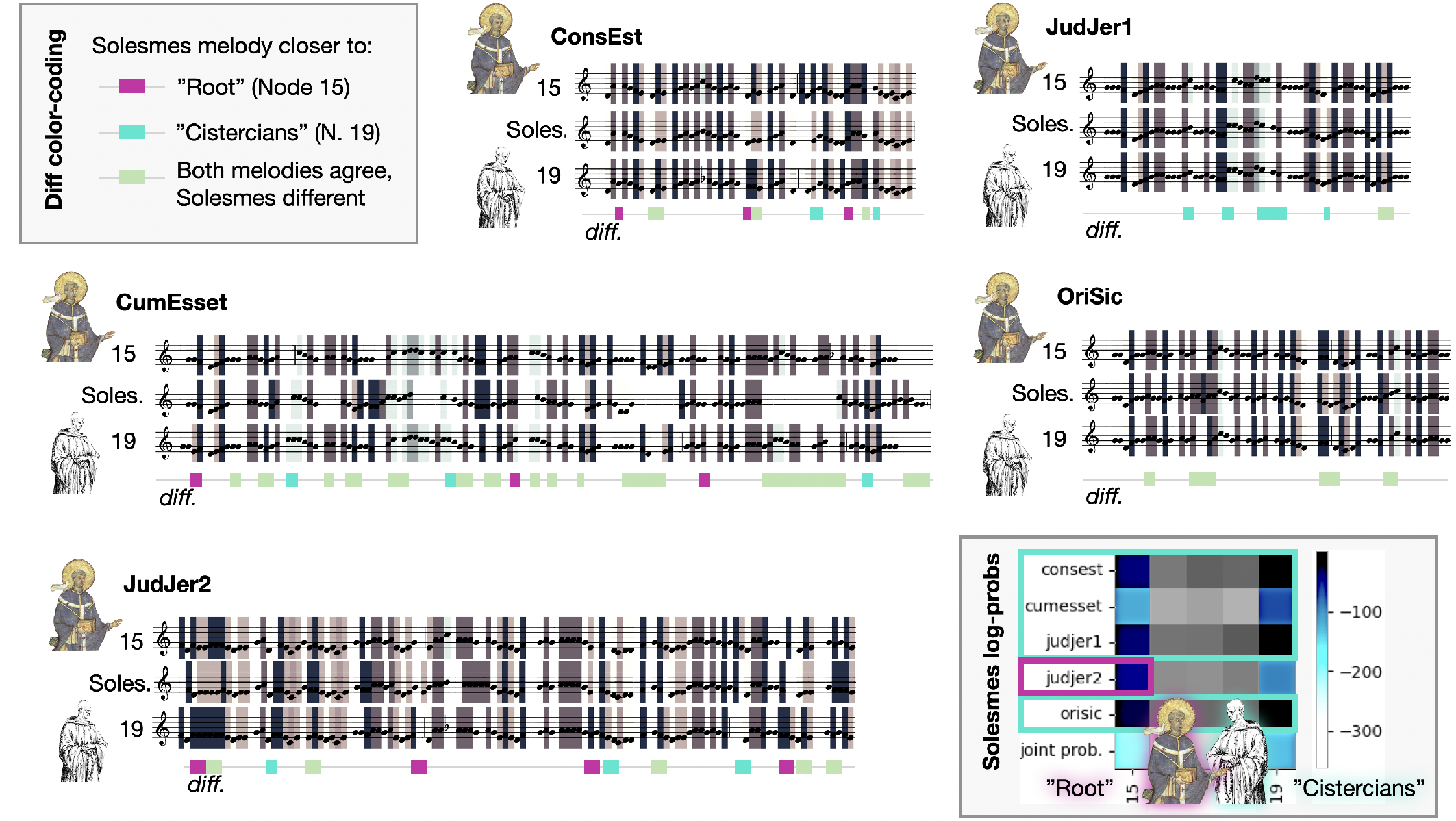
Analysis of the reconstructed melodies for the inferred root (node 15 of the DTE tree), the common ancestor of the later Cistercian sources (node 19 of the DTE tree), and the Solesmes version of the melodies. The log probability of observing the Solesmes melody at the Cistercian node is higher for each of the five melodies (the antiphon *Bethleem non es minima* is not in the Solesmes editions) except for the responsory Judaea et Jerusalem. Under each of the melody comparisons, we color-code the positions where Solesmes agrees with the root and not with the Cistercians (**purple**), positions where Solesmes agrees with the Cistercians rather than the root (**teal**), and positions where both inferred melodies agree but the Solesmes edition differs (**light green**), against the MAP estimates of the melodies obtained from ASR. Most of the differences are between the Solesmes edition and both of the inferred medieval melodies: thus one can observe where the Solesmes editorial process prioritized other criteria than fidelity to the earliest possible sources.

The melodies thus cannot be used to directly evaluate how well the root melody is reconstructed by ASR. Instead, observations can be made about the results of the Solesmes efforts: what was the extent to which these were “archaeological”, and what influence did “legitimate tradition” and “common practices of present-day liturgy” have? Here we borrow the term that Peter Wagner applied in his polemic with the critics of Editio Vaticana [59], while we do not necessarily agree with its original disparaging nature, it is not chosen without merit to represent that particular methodological approach. How does this recent layer of chant tradition relate to the medieval sources? These results serve to show how the inferred phylogeny can be used to provide insights complementary to philological methods, and perhaps also how the editorial practices of Solesmes evolved throughout the 20th and 21st centuries.

### 4.1 Future work

Further evaluation should be done on other data, such as the Officium Defunctorum study by Ottosen [65].

The method could then be applied to existing open problems in chant transmission, such as the issue of transmission of the “Germanic” melodic dialect [4, pp.116-117]. The fact that DTE predict a slightly earlier common ancestor for the east Frankish sources may be a part of the discussion, although more sources in the dataset are needed to try and narrow the time ranges down. Similarly, given an encoding of melodies from Old Roman sources, this method could provide additional empirical evidence of its chronology in relation to Gregorian chant, which is still an open problem [13, p. 466].

Another suggestion would be examining vernacular chant traditions, such as the early 15th century Jistebnice Cantionale [66] and later utraquist liturgical books in Bohemia [67], or modern editions of Chinese vernacular chant [68, 69] or Korean chant [70].

Another open problem is early chant transmission in adiastematic sources. Here, phylogenies could utilize sequences of neumes and/or visual features of the notation, rather than the melodies themselves. MEI supports adiastematic neumes, but manual encoding is costly and few have the expertise. However, as opposed to finding statistical patterns through machine learning, the phylogenetic pipeline has shown usefulness already with less than 100 transcribed sequences, making it a cost-effective option. Furthermore, by placing an adiastematic source into the phylogenetic tree, one can perhaps use the inferred ancestral melody to help resolve ambiguities of pitch when they arise.

Further work on Solesmes melodies may also bring historical chant research in a closer relationship to current living practice, which is based on these editions. Specifically, the activities of the schola in Kiedrich (see https://www.kiedricher-chorbuben.de/index.php/musik/choral) in reviving the Germanic dialect (with a papal dispensation), in combination with the Neumz project recorded at a French Benedictine abbey (see https://neumz.com/) can provide a stepping stone towards an ability to process the audio modality and start including features of performance practice in analyses. Using melody alone, not on genre-specific features such as liturgical positions, also enables tracing melodic evolution across repertoires. Can geographically defined melodic dialects show commonality with, e.g., regional folksong traditions? Or, can we perhaps even find traces of mostly lost pre-Gregorian repertoires such as Gallican chant or the chant of the British Isles?

Whatever the future findings, we are excited for the new options that this phylogenetic pipeline brings to the field of not only computational chant scholarship.

## 5 Acknowledgments

GAB was funded through a postdoctoral fellowship from FAPESP (process #23/07838-1). GAB thanks M. dos Reis S. Álvarez-Carretero, M. Panchaksaram, Z. Yang, A. Stamatakis, A. Ciliou, J. Thorne, B. Redelings, B. Bosseau, R. Warnock, A. Leaché, B. Rannalla, M. May, G. Tiley, and C. Solís-Lemus for fruitful discussions on computational phylogenetics. JH and KHM were funded by the Genome of Melody grant, supported by the Cultural Evolution Society Transformation Fund; underwritten by the John Templeton Foundation, Grant #61913. JH and KHM thank H. Vlhová-Wörner, D. Eben, C. Atkinson, T. Eipert, F. Moss, J. Ciglbauer, S. Street, and S. Škoviera for helpful discussions on the properties of Gregorian chant and its development. The opinions expressed in this publication are those of the author(s) and do not necessarily reflect the views of the John Templeton Foundation.

## References

[1] Dom Raphael Molitor. Our Position. A Word in Reference to the Plain Chant Question. F. Pustet, Printer to the Holy See, 1904.

[2] Peter Wagner. Einführung in die gregorianischen Melodien: t. Gregorianische Formenlehre; eine choralische Stilkunde, volume 3. Breitkopf & Härtel, 1921.

[3] Peter Wagner. Germanisches und Romanisches in Frühmittelalterlichen Kirchengesang. Verlag Breitkopf & Härtel, 1925.

[4] Alexander Blachly. Some observations on the” germanic” plainchant tradition. Current Musicology, (45-47):85–117, 1990.

[5] Debra Lacoste. The Cantus Database and Cantus Index Network. In The Oxford Handbook of Music and Corpus Studies. Oxford University Press, 2022.

[6] Johann-Mattis List, Shijulal Nelson-Sathi, Hans Geisler, and William Martin. Networks of lexical borrowing and lateral gene transfer in language and genome evolution. Bioessays, 36(2):141–150, 2014.

[7] David Hiley. Western plainchant: a handbook. Clarendon Press, Oxford, United Kingdom, 1993.

[8] Mary Carruthers and Jan M Ziolkowski. The medieval craft of memory: An anthology of texts and pictures. University of Pennsylvania Press, 2002.

[9] Anna Maria Busse Berger. Medieval music and the art of memory. Univ of California Press, 2005.

[10] Luisa Nardini. “ god is witness”: Dictation and the copying of chants in medieval monasteries. Musica disciplina, 57:51–79, 2012.

[11] Leo Treitler. The” unwritten” and” written transmission” of medieval chant and the start-up of musical notation. The Journal of Musicology, 10(2):131–191, 1992.

[12] Jan Hajič jr., Gustavo A. Ballen, Klára Hedvika Mühlová, and Hana Vlhová-Wörner. Towards Building a Phylogeny of Gregorian Chant Melodies. In Proceedings of the 24th International Society for Music Information Retrieval Conference, pages 571–578. ISMIR, December 2023.

[13] Helmut Hucke. Toward a new historical view of gregorian chant. Journal of the American Musicological Society, 33(3):437–467, 1980.

[14] Leo Treitler. The early history of music writing in the west. Journal of the American Musicological Society, 35(2):237–279, 1982.

[15] David G Hughes. Evidence for the traditional view of the transmission of gregorian chant. Journal of the American Musicological Society, 40(3):377–404, 1987.

[16] Patrick E Savage. Cultural evolution of music. Palgrave Communications, 5(1):1–12, 2019.

[17] Simon J. Greenhill. Language Phylogenies: Modelling the Evolution of Language. In The Oxford Handbook of Cultural Evolution. Oxford University Press, 2023.

[18] Roope O Kaaronen, Allison K Henrich, Mikael A Manninen, Matthew J Walsh, Isobel Wisher, Jussi T Eronen, and Felix Riede. The ties that bind: Computational, cross-cultural analyses of knots reveal their cultural evolutionary history and significance, Jun 2024.

[19] David N Matzig, Ben Marwick, Felix Riede, and Rachel C M Warnock. A macroevolutionary analysis of european late upper palaeolithic stone tool shape using a bayesian phylodynamic framework. Royal Society Open Science, 11(8):240321, 2024.

[20] Sylvie Le Bomin, Guillaume Lecointre, and Evelyne Heyer. The evolution of musical diversity: The key role of vertical transmission. PLOS ONE, 11(3):e0151570, March 2016.

[21] Patrick E. Savage, Sam Passmore, Gakuto Chiba, Thomas E. Currie, Haruo Suzuki, and Quentin D. Atkinson. Sequence alignment of folk song melodies reveals cross-cultural regularities of musical evolution. Current Biology, 32(6):1395–1402.e8, 2022.

[22] Mason Youngblood, Karim Baraghith, and Patrick E Savage. Phylogenetic reconstruction of the cultural evolution of electronic music via dynamic community detection (1975–1999). Evolution and Human Behavior, 42(6):573–582, 2021.

[23] Jonathan Warrell, Leonidas Salichos, Michael Gancz, and Mark B Gerstein. Latent evolutionary signatures: a general framework for analysing music and cultural evolution. Journal of the Royal Society Interface, 21(212):20230647, 2024.

[24] Alan Lomax. Factors of musical style. In Theory & practice: Essays presented to gene weltfish, pages 29–58. Mouton The Hague, 1980.

[25] P. Jeffery. Re-Envisioning Past Musical Cultures: Ethnomusicology in the Study of Gregorian Chant. Chicago Studies in Ethnomusicology. University of Chicago Press, 1992.

[26] Bas Cornelissen, Willem H Zuidema, John Ashley Burgoyne, et al. Mode classification and natural units in plainchant. In Proceedings of the 21st Int. Society for Music Information Retrieval Conf., pages 869–875, Montreal, Canada, 2020.

[27] Bas Cornelissen, Willem Zuidema, and John Ashley Burgoyne. Studying large plainchant corpora using chant21. In 7th International Conference on Digital Libraries for Musicology, pages 40–44, 2020.

[28] Kate Helsen, Mark Daley, and Jake Schindler. The sticky riff: Quantifying the melodic identities of medieval modes. Empirical Musicology Review, 16(2):312–325, 2021.

[29] Vojtěch Lanz. Unsupervised segmentation of gregorian chant melodies for exploring chant modality. 2023.

[30] Vojtěch Lanz and Jan Hajič. Text boundaries do not provide a better segmentation of gregorian antiphons. In Proceedings of the 10th International Conference on Digital Libraries for Musicology, pages 72–76, 2023.

[31] Walter Howard Frere. Antiphonale Sarisburiense: a reproduction in facsimile of a manuscript of the 13th century, with a dissertation and analytical index. Gregg Press Limited, 1901.

[32] Kate Helsen. The use of melodic formulas in responsories: constancy and variability in the manuscript tradition. Plainsong & Medieval Music, 18(1):61–76, 2009.

[33] John Glasenapp. To Pray without Ceasing: A Diachronic History of Cistercian Chant in the Beaupré Antiphoner (Baltimore, Walters Art Museum, W. 759–762). Columbia University, 2020.

[34] A Scarcez. L’Antiphonaire 12 A-B de Westmalle Dans L’Histoire Du Chant Cistercien Au Xiie Siecle. Brepols, Tournhout, Belgium, December 2011.

[35] Z Yang and B Rannala. Bayesian phylogenetic inference using DNA sequences: a Markov Chain Monte Carlo Method. Molecular Biology and Evolution, 14(7):717–724, 07 1997.

[36] Fredrik Ronquist and John P. Huelsenbeck. MrBayes 3: Bayesian phylogenetic inference under mixed models. Bioinformatics, 19(12):1572–1574, 08 2003.

[37] Joseph Felsenstein. Maximum-likelihood estimation of evolutionary trees from continuous characters. American Journal of Human Genetics, 25(5):471–492, September 1973.

[38] Jeffrey L. Thorne and Hirohisa Kishino. Divergence Time and Evolutionary Rate Estimation with Multilocus Data. Systematic Biology, 51(5):689–702, 2002.

[39] Bruce Rannala and Ziheng Yang. Bayes Estimation of Species Divergence Times and Ancestral Population Sizes Using DNA Sequences From Multiple Loci. Genetics, 164(4):1645–1656, 2003.

[40] Alexei J. Drummond and Andrew Rambaut. Beast: Bayesian evolutionary analysis by sampling trees. BMC Evolutionary Biology, 7(214):1–8, 2007.

[41] Joseph W Brown, Joseph F Walker, and Stephen A Smith. Phyx: phylogenetic tools for unix. Bioinformatics, 33(12):1886–1888, 2017.

[42] Wangang Xie, Paul O Lewis, Yu Fan, Lynn Kuo, and Ming-Hui Chen. Improving marginal likelihood estimation for Bayesian phylogenetic model selection. Systematic biology, 60(2):150–160, 2011.

[43] Emmanuel Paradis and Klaus Schliep. ape 5.0: an environment for modern phylogenetics and evolutionary analyses in R. Bioinformatics, 35:526–528, 2019.

[44] Liam J. Revell. phytools 2.0: an updated R ecosystem for phylogenetic comparative methods (and other things). PeerJ, 12:e16505, 2024.

[45] R Core Team. R: A Language and Environment for Statistical Computing. R Foundation for Statistical Computing, Vienna, Austria, 2024.

[46] Z. Yang. Molecular Evolution: A Statistical Approach. Oxford University Press, 2014.

[47] Tracy A. Heath, John P. Huelsenbeck, and Tanja Stadler. The fossilized birth–death process for coherent calibration of divergence-time estimates. Proceedings of the National Academy of Sciences, 111(29):E2957– E2966, 2014.

[48] R.C.M. Warnock and A.M. Wright. Understanding the Tripartite Approach to Bayesian Divergence Time Estimation. Elements of Paleontology. Cambridge University Press, 2020.

[49] Paul O Lewis. A likelihood approach to estimating phylogeny from discrete morphological character data. Systematic biology, 50(6):913–925, 2001.

[50] Thomas Lepage, David Bryant, Hervé Philippe and Nicolas Lartillot. A General Comparison of Relaxed Molecular Clock Models. Molecular Biology and Evolution, 24(12):2669–2680, 2007.

[51] Gautam Altekar, Sandhya Dwarkadas, John P. Huelsenbeck, and Fredrik Ronquist. Parallel Metropolis coupled Markov chain Monte Carlo for Bayesian phylogenetic inference. Bioinformatics, 20(3):407–415, 2004.

[52] A Rambaut and AJ Drummond. FigTree version 1.3.1 [computer program], 2009.

[53] Luke J. Harmon. Phylogenetic Comparative Methods: Learning from Trees. Open textbook library. CreateSpace Independent Publishing Platform, 2019.

[54] Liam J. Revell and Luke J. Harmon. Phylogenetic Comparative Methods in R. Princeton University Press, 2022.

[55] Microsoft Corporation and Steve Weston. doParallel: Foreach Parallel Adaptor for the ‘parallel’ Package, 2022. R package version 1.0.17.

[56] Microsoft and Steve Weston. foreach: Provides Foreach Looping Construct, 2022. R package version 1.5.2.

[57] Kazutaka Katoh and Daron M. Standley. MAFFT Multiple Sequence Alignment Software Version 7: Improvements in Performance and Usability. Molecular Biology and Evolution, 30(4):772–780, 01 2013.

[58] T. A. Burge. The vatican edition of the kyriale and its many critics. In Irish Ecclesiastical Record, pages 324–345, 1906.

[59] Peter Wagner. The attack on the vatican edition: a rejoinder. Caecilia: A review of Catholic Curch Music, 87(1):10–44, orig. 1907, transl. 1960.

[60] Julie Kerr. An Essay on Cistercian Liturgy. Cistercians in Yorkshire, University of Sheffield, 2012.

[61] Maria Mairold. Die datierten Handschriften in der Steiermark außerhalb der Universitätsbibliothek Graz bis zum Jahre 1600, volume 7 of Katalog der datierten Handschriften in lateinischer Schrift in Österreich. Österreichische Akademie der Wissenschaften, Wien, 1988.

[62] Pierre Combe, Theodore N. Marier, and William Skinner. The Restoration of Gregorian Chant: Solesmes and the Vatican Edition. Catholic University of America Press, 2003.

[63] Henry Bewerunge. The vatican edition of plain chant. In Irish Ecclesiastical Record, pages 44–63, 1906.

[64] Katharine Ellis. The Politics of Plainchant in fin-de-siècle France, volume 20 of ROYAL MUSICAL ASSOCIATION MONOGRAPHS. Routledge, 2013.

[65] Knud Ottosen. The responsories and versicles of the Latin office of the dead. BoD–Books on Demand, GmbH, 2007.

[66] David Holeton, Jaroslav Kolár, Anežka Vidmanová, and Hana Vlhová-Wörner. Jistebnice kancional: MS. Prague, national museum library II C 7. L. Marek, 2005.

[67] David R Holeton. The evolution of utraquist liturgy: a precursor of western liturgical reform. Studia liturgica, 25(1):51–67, 1995.

[68] Lionel Li-Xing Hong. Catholic music in seventeenth and eighteenth century china : a study from a liturgical perspective. In Atti del Congresso internazionale di musica sacra : in occasione del centenario di fondazione del PIMS, Roma, 26 maggio - 1 giugno 2011, pages 1231–1245. Pontificio Instituto di Musica Sacra, 2013.

[69] Lionel Li-Xing Hong. Practice of catholic liturgical music in taiwan today : an overview. In Atti del Congresso internazionale di musica sacra : in occasione del centenario di fondazione del PIMS, Roma, 26 maggio - 1 giugno 2011, pages 1373–1378. Pontificio Instituto di Musica Sacra, 2013.

[70] Eun Young Cho, Hayoung Wong, and Zong Woo Geem. The liturgical usage of translated gregorian chant in the korean catholic church. Religions, 12(12), 2021.

